# Diffuse Reflectance Spectroscopy reveals heat stress-induced changes in hemoglobin profile in chicken breast

**DOI:** 10.1101/741520

**Authors:** Sina Dadgar, Liz Greene, Ahmed Dhamad, Barbara Mallmann, Sami Dridi, Narasimhan Rajaram

## Abstract

Global rise in incidence of woody breast (WB) syndrome imposes a significant economic burden on the poultry industry. The increase in WB is due to the large increase in the weight of chickens these days within a very short period. An early determination of WB can significantly reduce losses to the poultry industry. Diffuse reflectance spectroscopy provides a noninvasive and rapid method to interrogate tissue function. The sensitivity of DRS to the distinct absorption spectra of oxygenated and deoxygenated hemoglobin allows accurate quantification of average hemoglobin concentration and vascular oxygenation within the sampled tissue. In this study, we used diffuse reflectance spectroscopy to monitor breast hemoglobin concentration (THb) and vascular oxygen saturation (sO_2_) of 16 chickens that were exposed to heat stress (HS). HS is an important cause of WB myopathy in chickens. Animals were exposed to heat-stress (HS) and optical data were acquired at three time points: at baseline prior to heat stress, 2 days, and 21 days after initiation of HS. Our results show that animals from control and HS groups had a steady decay in optically derived breast hemoglobin concentration consistent with independent i-STAT measurements made on blood sampled from the femoral artery and could provide a noninvasive technology for monitoring tissue function in the poultry industry.

## 1 Introduction

Poultry meat is a mass consumer product and one of the main food sources worldwide for billions of people. In 2006, US was the largest consumer of poultry meat with average consumption rate of 54 kg/capita/year [1]. Boneless breast meat is a popular choice for consumers and high breast-yielding strains of broilers are adopted to meet this fast-growing demand. Today chickens and turkeys are marketed in twice the body weight in shorter period of time compared to 50 years ago [2]. This increase in body weight within such short periods of time can lead to various meat quality problems including high incidence of metabolic disorders such as woody breast (WB) myopathy [3]. Emerging in global scale, WB is reported to have extreme palpable stiffness of breast muscle which severely affects meat appearance with bulge-out and pale color [4]. WB can adversely affect consumer acceptance of poultry meat which can result in huge economic loss to the industry. Optical methods, such as near infrared (NIR) spectroscopy have been demonstrated to correctly differentiate WB fillets from normal fillets based on their water and protein content [5,6] because WB is known to contain more loosely water bound and lower protein content compared to normal breast muscle [7]. Although NIR spectroscopy allows for rapid identification of WB fillets in production line, they are not capable of identifying animals susceptible of formation of WB. Although several transcriptomic and proteomic studies relate WB myopathy to localized muscular hypoxia [8] and oxidative stress [9], unfortunately no method yet reliably determines the likelihood of WB formation in chickens at earlier time points. Thus, there remains a critical unmet need to engineer new predictive biomarkers of WB formation. Such a biomarker can identify animals susceptible of WB formation in advance to avoid further investment in those animals. In order to fill this gap, we propose to use diffuse reflectance spectroscopy (DRS) to monitor the hemoglobin concentration and vascular oxygen saturation of the chicken breast. DRS is an optical fiber-based technique that uses ‘source’ optical fibers to deliver low-power non-ionizing broadband white light to the tissue surface and ‘detector’ optical fibers to collect the diffusely reflected light from tissue. The source and detector fibers are typically separated from each other, with the source-detector separation distance determining the sampling depth within tissue. This diffusely reflected light carries the spectral signatures of tissue components that interacted with the light. These ‘interactions’ consist of 1. Elastic scattering (no loss of energy but a change in direction) from cell nuclei, mitochondria and collagen, and 2. Absorption, primarily by oxygenated and deoxygenated hemoglobin in blood vessels. Depending on tissue type, other major absorbers include melanin (skin) and beta-carotene (breast). By analyzing the diffusely reflected light from tissue using analytical or quantitative models of light-tissue interaction, it is possible to determine the scattering and absorption properties of interrogated tissue, and extract meaningful information regarding tissue scattering, hemoglobin concentration, and vascular oxygen saturation. Specifically, the distinct absorption spectra of oxygenated (HbO_2_) and deoxygenated hemoglobin (dHb) allow the quantification of total hemoglobin concentration (THb = HbO_2_ + dHb) and vascular oxygen saturation (sO_2_ = HbO_2_/THb) in the tissue sampling volume. Previous work by us and others have demonstrated that vascular oxygenation measured using DRS is correlated with immunohistochemical measures of hypoxic fraction [10] and microelectrode based measures of tissue oxygenation (pO_2_) [11,12]. One of the foremost applications of diffuse reflectance spectroscopy has been in the field of cancer as a potential complementary tool to histopathology. These applications have spanned early cancer detection in several organs including skin [13] and breast [14], identification of surgical margins [15], and monitoring response to therapy [16–18]. The underlying premise of these applications is that the major sources of tissue scattering and absorption are also molecules that undergo significant changes during disease progression or in response to external stimuli; therefore DRS can provide a noninvasive and nonionizing ‘optical biopsy’ of tissue. The simplicity of the tool also lends itself to potentially repeated measurements on the same tissue, thus providing a continuous time-lapse of changes in tissue rather than a single snapshot.

Given the sensitivity of DRS to changes in vascular oxygenation and the relevance of tissue hypoxia in chicken breast pathology, we sought to determine the effects of heat stress (HS) in chickens using DRS. HS is devastating to poultry production due to its adverse effects on growth performance and mortality. The most prominent effect of HS is depression in feed intake and increase in core body temperature. To increase heat loss, birds divert blood to the periphery (skin) which results in a hypoxia-like state in the internal organs such as the breast muscle. We have used DRS to show that circulatory and breast muscle oxygen homeostasis is deregulated in chickens with WB myopathy compared to healthy counterparts (data not shown). In this study, we determined hemoglobin concentration in chickens exposed to heat stress (HS) and compared our measurements with the standard i-STAT device for measuring circulatory Hb levels. Our results show that aging leads to a decline in optically derived breast hemoglobin levels, a result that was confirmed by independent i-STAT measurements. Furthermore, longitudinal changes in optically derived Hb concentration were consistent with i-STAT measurements. Our data indicate that DRS measures of hemoglobin concentration could be a reliable, rapid, accurate, and non-invasive method to measure hematologic parameters in avian species.

## 2 Materials and Methods

### 2.1 Animal storage and handling

The studies with chickens were approved by the University of Arkansas Institutional Animal Care and Use Committee (IACUC Protocol #16084). 3-week-old chickens (n=16) were randomly divided into cyclic heat stress (HS, 35°C for 12 hours/day) and thermoneutral conditions (TN, 24°C) and were given *ad libitum* access to food and clean water. We collected optical data and blood samples at three time points: before exposure to heat-stress (pre-HS), 2 days after initiation of HS (Acute-HS), and 21 days after HS (Chronic-HS). First, 10 mL of blood sample were collected from femoral artery from each animal. This was followed by optical data acquisition from chicken breast while animals were held upside down. In addition, we determined the presence of WB by manual palpation as previously described in literature [19].

### 2.2 Diffuse reflectance spectroscopy

Our spectroscopic system consists of a halogen lamp (HL-2000, Ocean Optics, Dunedin, Florida) as light source and a USB spectrometer (Flame, Ocean Optics) for spectral light acquisition. The common end of a bifurcated probe was employed for delivery and collection of light with four illumination and five detector fibers located at a source-detector separation distance (SDSD) of 2.25 mm. We have confirmed that this probe samples light from a depth of approximately 1.8 mm [20]. Data acquisition was simplified by using a foot pedal controlled by a custom LabVIEW (National Instruments, Austin, Texas) software. Minimum of 15 spectral data in spectral range of 475 to 600 nm were collected from multiple sites from each chicken’s breast and averaged optical properties were used to represent that chicken. Prior to any optical measurement from animals, reflected light intensity of an 80% reflectance standard (SRS-80-010; Labsphere, North Sutton, New Hampshire) was acquired to calibrate for daily variations in light throughput.

### 2.3 Quantification of tissue optical properties

We used a LUT-based inverse model to fit the acquired data and extract wavelength-dependent absorption and scattering properties of tissue. The model has been described previously [21] and validated for a range of different source detector separations (SDSDs) [20]. To fit the model to the acquired optical spectra, we constrained scattering to follow a negative power-law dependence on wavelength [22]: 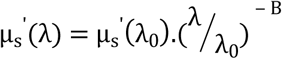 with *λ*_0_ =600 nm as a reference point where light absorption is minimum. We further assumed light absorption to be a linear sum of light absorbing chromophores, namely oxygenated and deoxygenated hemoglobin, as well absorption from skin. In the spectral range of 475-600 nm, we calculated µ_a_as: 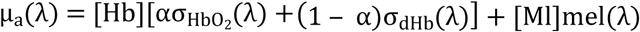, where [Hb] and [Ml] respectively are total hemoglobin concentration and skin absorption. α is oxygen saturation which represents the ratio of oxygenated (HbO_2_) to total hemoglobin concentration [Hb]. The fixed absorption parameters, extinction coefficients of oxygenated hemoglobin 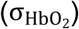, deoxygenated hemoglobin (σ_dHb_), and melanin (mel) were obtained from an online database [23]. Generation of LUT and data analysis was performed in MATLAB (Mathworks, Natick, Massachusetts).

### 2.4 i-STAT system

Circulatory hemoglobin concentration (Hb) were determined using i-STAT Alinity system (SN: 801128; software version JAMS 80.A.1/CLEW D36; Abaxis, Union City, CA) with the i-STAT CG8+ cartridge test (ABBT-03P77-25) according to manufacturer’s recommendation. i-STAT derived hemoglobin values have previously been validated in laying hens [24,25].

### 2.5 Statistical analysis

Repeated measures analysis of two-factor ANOVA was employed to determine statistically significant differences in Hb levels from both DRS and i-STAT readings between different groups. Time and treatment were respectively considered as within and between effects. Additionally, interactions between all effects were included in the analysis. Post-hoc Tukey HSD test were used to differentiate specific groups. Significant differences among slopes of regression lines were tested with Analysis of Covariance (ANCOVA). All statistical analyses were performed using JMP (ANOVA) and GraphPad Prism (ANCOVA).

## 3 Results

Absorption spectra of oxyegnated (HbO_2_ - solid line) and deoxygenated hemoglobin (dHb – dashed line) are illustrated in **figure 1A**. **Figure 1B** presents the measured diffuse reflectance spectra from representative animals in TN (orange circles) and HS (green circles) groups and their corresponding LUT fit (**black line**) at pre-HS time points. Based on LUT fits, we extracted absoprtion (**Fig 1C**) and scattering (**Fig 1B**) properties of the reflectance data. We noted absorption spectra belonging to HS have a slightly more pronounced Hb bands compared to that of TN group. We additionally observed similar magnitude of scattering in HS and TN groups at pre-HS time point.

**Figure 1.**
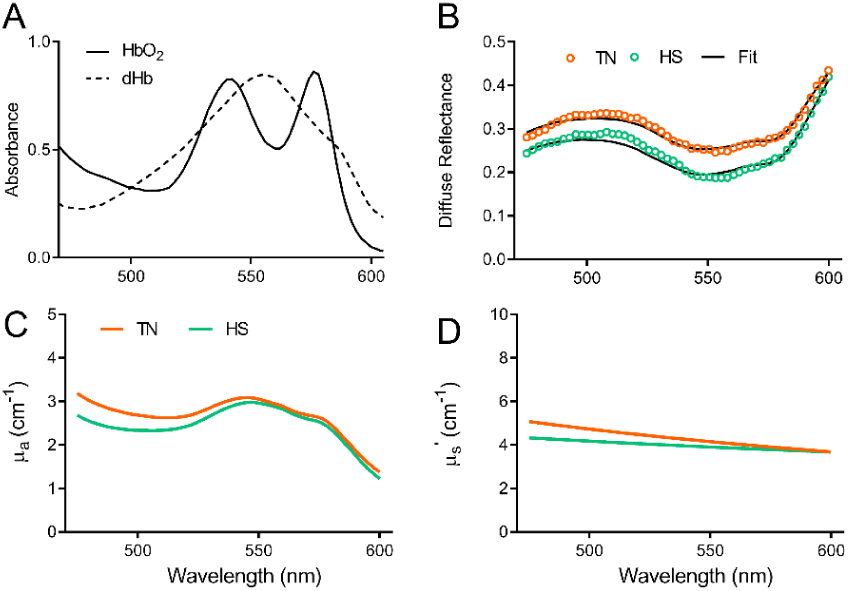
(A) Absorbance spectra of oxygenated and deoxygenated Hb. (B) Measured diffuse reflectance spectra of representative animals in TN (orange circles) and HS (red circles) and their corresponding LUT fit (black line). LUT extracted absorption (C) and scattering (D) coefficients.

To enable quantitative comparisons between the different groups, we determined the fit parameters derived from the absorption and scattering coefficients. **Figure 2** shows the temporal kinetics of the concentration of i-STAT-measured Hb (**Figure 2.A**), concentration of DRS-measured Hb (**Fig 2B**), concentration of dHb (**Figure 2C**), concentration of HbO_2_ (**Figure 2D**), oxygen saturation (**Fig 2E**), and mean reduced scattering coefficient (**Fig 2F**). For each treatment, the data are presented as pre-HS (**blue bars**), Acute-HS (**green bars**), and Chronic-HS (**red bars**) ± standard error of the mean (SEM). **Figures 2A & 2B** demonstrate a trend toward lower Hb concentration after HS, using both the i-STAT and DRS devices. While no significant differences between groups were found in i-STAT measurements of circulatory Hb, [Hb] measured in the chicken breast using DRS was significantly lower at the chronic HS time point compared with pre-HS and acute-HS. While the decrease in [Hb] was observed in both TN and HS groups over time, the source of this decrease appears to be different in both groups. Specifically, there was a significant decrease in [dHb] concentration in the chronic-HS TN group. On the other hand, the [Hb] decrease in the HS group appears to be driven primarily by a large, significant decrease in [HbO_2_] in the HS group. To further investigate these group-specific changes, we calculated the vascular oxygen saturation (sO_2_) as the ratio of HbO_2_/(dhb+HbO_2_) illustrated in **fig 2E**. We found statistically significant differences in sO_2_ between the acute and chronic time points of the HS group. No significant differences were found in the TN group. Finally, we found significant decreases in the mean reduced scattering coefficient between pre-HS and acute HS time points in both the TN and HS groups, followed by a significant increase at the chronic time point.

**Figure 2.**
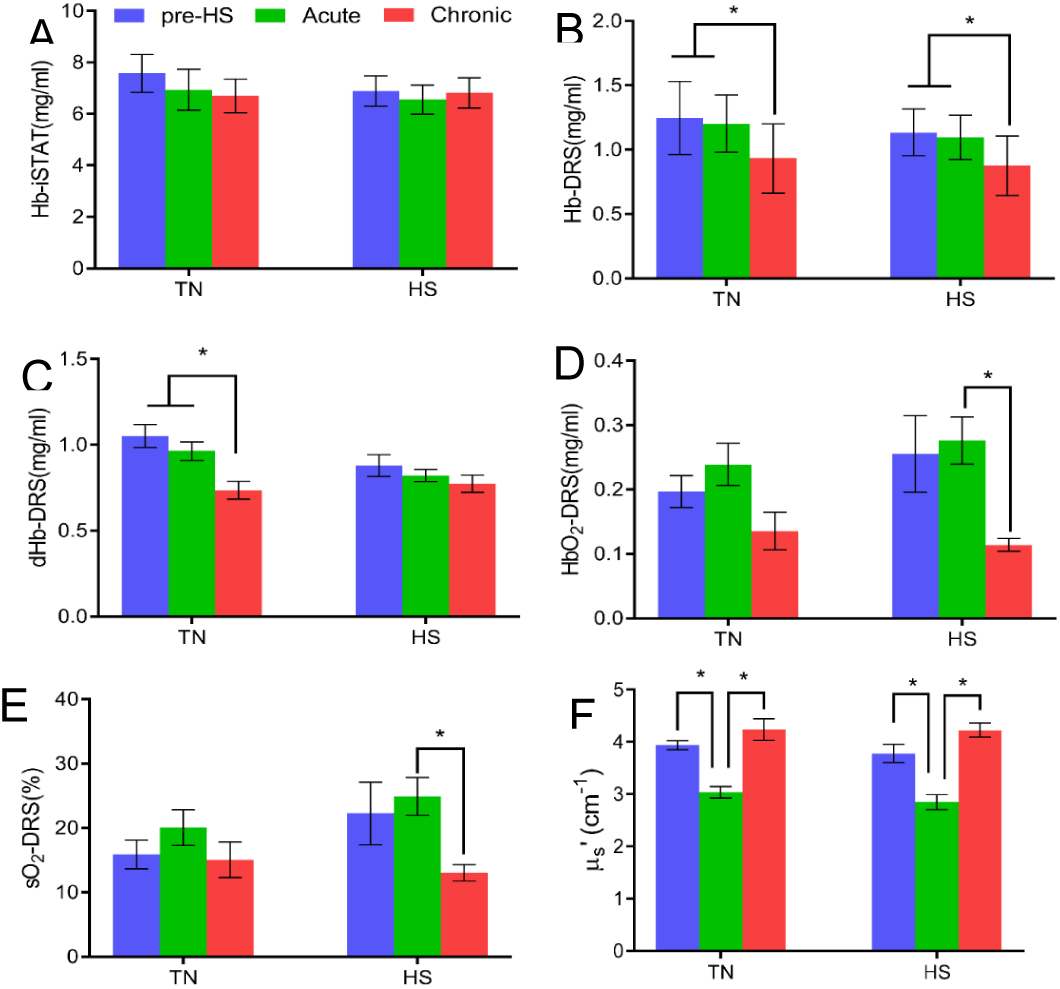
Circulatory hemoglobin concentration detected by i-STAT device (A). Optically derived parameters: Total hemoglobin concertation – Hb (B), deoxygenated hemoglobin – dHb (C), oxygenated hemoglobin – HbO_2_ (D), hemoglobin oxygen saturation – sO_2_ (E), and total scattering coefficient – µ_s_’ (F). Data shown as means ± SEM representing pre-HS (blue bars), Acute-HS (green bars), and Chronic-HS (red bars).

To determine whether the rates of change in optically measured Hb were similar to the i-STAT measurements, we compared the slope of regression lines over time (**Figure 3)**. There were no significant differences in the slope of the regression line in TN and HS groups. TN (P=0.17, **Fig 3A**), HS (p=0.65, **Fig 3B**). Note that hemoglobin concentration measured by i-STAT and DRS are illustrated in green and blue, respectively.

**Figure 3.**
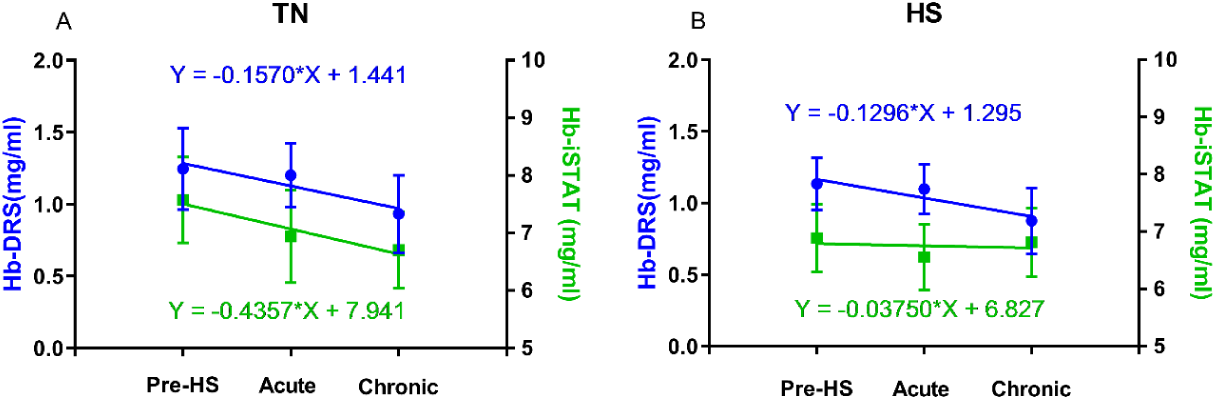
Longitudinal comparison of the slope of changes in Hemoglobin concentration in TN (A) and HS (B) groups detected with i-STAT (green) and DRS (blue). The equation of average line from each measurement are represented with their abovementioned colors.

In addition to collecting blood samples and optical spectra from each chicken, we determined the presence of WB by manual palpation. We observed that for both treatment groups, the fold change in the incidence rate of WB increased, with a higher rate observed in the TN group. (**Figure 4A**).

**Figure 4.**
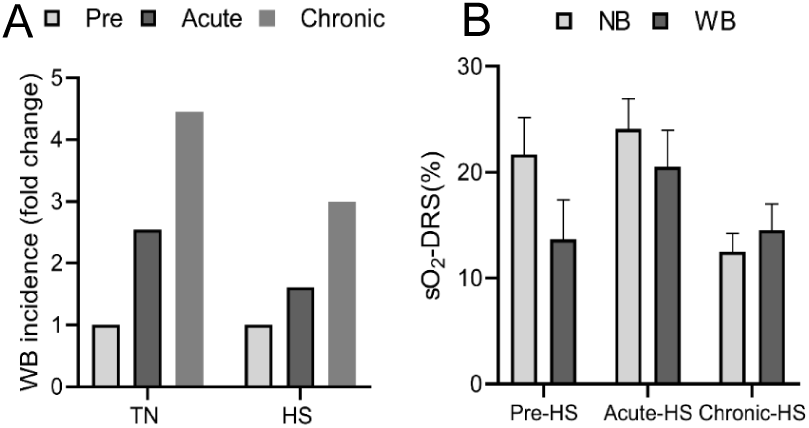
(A) Comparison of the fold change in incidence of WB in TN and HS groups at pre-HS, Acute-HS, and Chronic-HS time points. (B) Optically derived Hb readings in animals with normal breast (NB) and woody breast (WB) animals across time. Error bars are standard error of mean (SEM).

Since animals from both TN and HS groups formed WB, we sought to investigate whether vascular oxygen saturation (sO_2_) could differentiate animals with WB from those with normal breast. We classified chickens as normal breast (NB) or woody breast (WB) based on final assessment of the breast on Day 21. We found higher sO_2_ in animals with NB compared with animals with WB at pre-HS and acute-HS time points but not at the Chronic-HS time point. However, we failed to observe any statistically significant differences among the groups at any of the time points (**Figure 4B**).

## 4 Discussion

Previous studies have shown the importance of optically derived hemoglobin concentration in cancer research studies. For instance, hemoglobin concentration has been shown to be elevated in breast cancer patients due to angiogenesis [26,27]. In addition to that, near infrared (NIR) spectral tomography has been used to show that breast adipose and fibro-glandular tissues have significantly different contents of hemoglobin [28]. Mell et al. have shown that the volume of bone marrow is associated with hematologic toxicity of anal cancer patients treated with concurrent chemo-radiotherapy [29]. Crawford et al. have also associated the hemoglobin counts with quality of life improvements in anemic cancer patients treated with epotin alfa [30].

In this study, we used DRS to evaluate the effects of acute and chronic HS on hemoglobin profile in broiler chicken breast. We observed temporally significant decay in optically derived Hb concentration of animals exposed to TN and HS which were validated by i-STAT measurements. The similarity in slope of changes in hemoglobin detected by DRS and i-STAT indicate that aging can lead to a decay in both circulatory and breast hemoglobin concentration disregarding environmental conditions. The decrease in the circulatory and breast muscle hemoglobin concentration suggests a possible presence of circulatory (anemic) hypoxia which leads to diminished supply of oxygen to breast muscle [31](**Figure 2. E**). While sO_2_ in TN group remained relatively the same, oxygen saturation in HS group had a significant decrease after a sudden increase in Acute-HS time point which is due to significant decrease in the content of oxygenated hemoglobin [HbO_2_]. This significant decrease in sO_2_ value in chronic time-point of HS groups is negatively correlated (r^2^=-0.16) with palpation scores of WB incidence, however the correlation was not statistically significant (data not shown).

We observed an increase of light scattering in chronic-HS time points of both TN and HS groups (**Figure 2. F**). Chickens with WB myopathy have significantly higher levels of fibrosis, collagen, and necrosis [9], all of which can greatly contribute to light scattering [32]. However, we do not understand the etiology of reduction of light scattering at the acute-HS time point.

To examine how WB incidence changes over time and its association with TN and HS treatments, we determined the presence of WB by manual palpation. We observed that a greater increase in WB incidence in the TN group over time compared with the HS group (**Figure 4. A**). HS is known to lead to hyperthermia, which can cause a reduction in animal feed intake to avoid further diet-induced thermogenesis (DIT) [33]. This reduction in feed intake and body weight likely explains why HS chickens had a lower incidence of WB.

We showed that optically derived oxygen saturation is higher in animals with healthy breast (NM) compared to animals with WB. This difference was observed at pre-HS and acute-HS time-points. However, the opposite was observed 3 weeks after initiation of the study. More studies with larger sample size are required to establish the sensitivity of such optically derived parameters.

This study shows that diffuse reflectance spectroscopy has the potential to dynamically monitor changes in physiology and morphology of chicken breast. Further research with larger sample size with possibility of animal euthanasia at acute-HS time point can shine more light into HS, WB, and their relationship and can aid in early determination of WB formation. Unlike the i-STAT, DRS provides a method to measure vascular changes directly from the breast. An accurate early determination, which is not possible by manual palpation, can aid in developing strategies to treat WB. The low cost and easy implantation of DRS setup makes it an ideal screening tool for longitudinal monitoring of WB formation.

## ACKNOWLEDGEMENTS

This work was supported by a grant from The Arkansas Bioscience Institute (to S.D.)

